# Psychological trauma increases blood pressure sensitivity to angiotensin II via T-lymphocytes independent of psychopathology

**DOI:** 10.1101/2025.08.27.672648

**Authors:** Adam J. Case, Tamara Natour, Lauren J. Pitts, Tatlock H. Lauten, Emily C. Reed, Cassandra M. Moshfegh, Safwan K. Elkhatib

**Author notes:** **Corresponding author**: Adam J. Case, PhD Associate Professor Department of Psychiatry and Behavioral Sciences Department of Medical Physiology 8447 John Sharp Pkwy MREB2 3414 Bryan, TX 77807. **Conflict of Interest Statement:** The authors have declared that no conflict of interest exists.

## Abstract

Exposure to traumatic stress can lead to psychopathology, including post-traumatic stress disorder (PTSD), but may also cause inflammation and cardiovascular dysfunction. Clinical evidence suggests that exposure to traumatic stress, independent of psychopathology development, may be sufficient to induce pathophysiological sequelae, but this has not been thoroughly investigated. Using a model of repeated social defeat stress (RSDS), we explored links between the behavioral and physiological consequences of this paradigm. RSDS was sufficiently able to elevate systemic inflammation in both male and female mice, with no observed sex differences. RSDS also induced a heightened blood pressure sensitization response to low dose exogenous angiotensin II (AngII), suggesting the model was also sufficient in generating cardiovascular pathology. Interestingly, the RSDS-induced sensitization to AngII was completely abrogated in mice lacking T-lymphocytes (i.e., Rag2^-/-^ mice). Of note, Rag2^-/-^ mice demonstrated similar pro-social and anxiety-like behavior to wild-type mice, inferring that 1) differences in these behavioral outcomes do not explain the loss of RSDS-induced AngII sensitization in Rag2^-/-^ mice and 2) T-lymphocytes do not appear to impact these specific RSDS-induced behaviors. Indeed, intra-animal correlations demonstrate a tight association between the inflammatory and cardiovascular consequences post-RSDS, but no associations between these parameters with behavior. Together, our data suggest that exposure to traumatic stress, independent of psychopathology, is sufficient to induce pathophysiology. These findings have significant clinical implications, specifically for individuals who have experienced traumatic stress without the development of psychopathology, as this overlooked population may have similar risks of developing somatic diseases.

## 1 Introduction

It is estimated that 70-90% adults in the United States will have experienced one or more traumatic stressors, as defined by the Diagnostic and Statistical Manual, in their lifetime (1-3). The most common psychopathology to develop from these traumatic stressors is post-traumatic stress disorder (PTSD), which arises in approximately 10-40% of those exposed to a traumatic stressor (2, 4-6). This striking statistic suggests that >50% of individuals exposed to a traumatic stressor never develop PTSD (or possibly any other psychopathology), which would classify these individuals as more “resilient” to those who develop behavioral pathology. Indeed, a significant amount of research over the recent decades in both humans and animals has been devoted to identifying the underlying mechanisms that cause resiliency and susceptibility to psychopathology after traumatic stress (7-13). However, given that psychopathology is currently diagnosed exclusively by patient reported or observed behavioral changes (14), by definition, the characterization of resiliency and susceptibility has been defined only by the behavioral spectrum. In doing so, the impact of traumatic stressors on physiological changes throughout the body is often overlooked, not recorded, and it remains unknown if these somatic changes are coupled in any way to behavioral pathology.

A significant amount of research has demonstrated that PTSD increases the risk of developing comorbid pathologies such cardiovascular diseases (15-20). While many PTSD patients partake in activities that independently increase their risk for cardiovascular disease (*i.e.,* tobacco use, poor diet, and alcohol consumption), epidemiologic assessment has unequivocally demonstrated these confounding variables do not explain the increased risk for cardiovascular diseases in this patient population (21-24). This indicates a pathophysiological change is occurring in these patients with PTSD that predisposes them to altered cardiovascular function. However, by focusing specifically on PTSD, most of these studies have overlooked the majority population of individuals who experienced traumatic stress but did not develop overt symptoms to be classified with PTSD. The concept that psychological trauma alone, independent of the development of PTSD, is the underlying cause for the development of secondary pathophysiology is recently (albeit slowly) gaining traction. For example, both psychological trauma and PTSD share known biological effects that elevate the risk for inflammatory disorders, such as elevated sympathetic nervous system activity, immune dysregulation, redox imbalance, mitochondrial dysfunction, and alterations in the renin-angiotensin system (25). Given this, it is possible that the reason PTSD induces a high risk for inflammatory comorbidities is simply due to the underlying exposure to a traumatic stressor, as opposed to the development of behavioral abnormalities characteristic of PTSD. Therefore, studies specifically exploring outcomes and relationships between behavior and pathophysiology following psychological trauma are highly needed to address this hypothesis.

Immune cells, particularly T-lymphocytes, are overly sensitive to the psychobiological changes of traumatic stress, and T-lymphocytes are primary contributors to cardiovascular diseases including hypertension (26-29). For example, patients with PTSD have decreased numbers of naïve and regulatory (anti-inflammatory) T-lymphocytes with concurrent increases in pro-inflammatory memory T-lymphocytes (30, 31). Furthermore, circulating cytokines produced from T-lymphocytes, including interleukin 17A (IL-17A), are also elevated with PTSD (32-35). IL-17A is highly significant, as it has been shown to directly contribute to the pathogenesis of cardiovascular diseases including hypertension (36-41). However, studies investigating if traumatic stress (with or without behavioral changes) leads to cardiovascular disease via T-lymphocyte dysregulation are non-existent.

Herein, using an adapted novel model of chronic and severe traumatic stress in both male and female mice (i.e., repeated social defeat stress, RSDS), we investigated the impact of psychological trauma on the development of cardiovascular dysfunction, particularly hypertension. We found that while RSDS significantly elevated blood pressure during the active stress paradigm, blood pressure returned to unstressed control levels immediately after the traumatic stressor was removed. However, RSDS elevated sensitization to a mild cardiovascular challenge of low dose angiotensin II (AngII), where previously stressed animals demonstrated significantly higher blood pressure response to the AngII compared to unstressed controls. Moreover, this blood pressure sensitization to AngII appeared to be dependent upon IL-17A and T-lymphocytes. Importantly, while RSDS-induced blood pressure sensitization appeared tightly linked with systemic IL-17A levels, it was not linked to two major behavioral outputs of the RSDS paradigm (i.e., pro-social and anxiety-like behavior).

Together, our findings demonstrate that exposure to traumatic stress appears to lead to enhanced sensitization to cardiovascular dysfunction via immune dysregulation, and that these pathophysiological consequences of traumatic stress are not coupled with psychopathology. These results may have significant implications regarding the treatment of psychological trauma, as they suggest the absence of chronic behavioral pathology does not rule out potential for life-threatening somatic diseases as a result of the trauma alone.

## 2 Methods

### Mice

Wild-type C57BL/6J (#000664; shorthand WT), Rag2 knock-out mice (#008449; shorthand Rag2^-/-^), and estrogen receptor 1 alpha cre (#017913; shorthand Esr1-cre) mice were obtained from Jackson Laboratories (Bar Harbor, ME, USA). CD1 mice were purchased from Charles River (#022, Wilmington, MA, USA). Esr1-cre mice were backcrossed to CD1 mice >10 generations to create a congenic CD1 background strain. All mice were bred in house to eliminate shipping stress and microbiome shifts, as well as co-housed with their littermates (≤5 mice per cage) prior to the start of experimentation to eliminate social isolation stress. Mice were housed with standard pine chip bedding, paper nesting material, and given access to standard chow (#8604 Teklad rodent diet, Inotiv, West Lafayette, IN, USA) and water ad libitum. Male and female experimental mice between the ages of 8-12 weeks were utilized in all experiments. Experimental mice were randomized, and when possible, experimenters were blinded to the respective cohorts until the completion of the study. Mice were sacrificed by pentobarbital overdose (150mg/kg, Fatal Plus, Vortech Pharmaceuticals, Dearborn, MI, USA) administered intraperitoneally. All mice were sacrificed between 7:00 and 9:00 Central Time to eliminate circadian rhythm effects on T-lymphocyte function. All procedures were reviewed and approved by Texas A&M University Institutional Animal Care and Use Committee.

### RSDS

To overcome the limitation of using males only in RSDS, we have adapted a novel model of RSDS that utilizes chemogenetically altered aggressive mice that indiscriminately show aggression to both male and female experimental mice (42, 43). Briefly, male Esr1-cre mice are injected with Designer Receptors Exclusively Activated by Designer Drugs (DREADDs) viral constructs into the aggression center of the ventromedial hypothalamus. Administration of systemic clozapine-N-oxide (CNO) allows for temporal regulation of the DREADD constructs, which makes the mice hyperaggressive. For RSDS, experimental mice were placed in the home cage of a male Esr1-cre mouse (pre-injected with CNO) to induce a physical confrontation and fear induction for 1 minute given the highly aggressive nature of the chemogenetically-modified male Esr1-cre mice. Following the aggressive interactions, a clear perforated divider was placed into the cage and both mice remained co-housed for 24 hours. This process was repeated with a new aggressive Esr1-cre mouse daily for 10 consecutive days. Control mice were pair housed in the absence of an Esr1-cre mouse for the duration of the protocol. Mice were continuously monitored for visual wounding or lameness; None of the mice utilized herein met the threshold for exclusion (wounds >1 cm or any lameness).

### Behavior

For this study, pro-social and anxiety-like behaviors were assessed on Day 1 of the recovery period after RSDS. First, pro-social behaviors were evaluated using the social interaction test as previously described (44). Briefly, experimental mice were allowed to explore a standard open field chamber with an empty mesh cage at one end for 2.5 minutes. Following this, mice were tested again in the open field chamber in the presence of a novel aggressive Esr1-cre mouse in the mesh cage for an additional 2.5 minutes. Sessions were recorded and digitally analyzed by blinded reviewers using Noldus Ethovision XT 13 software (Leesburg, VA, USA). The social interaction zone ratio was calculated by dividing the time the mouse spent in the interaction zone close to the mesh cage in the presence versus the absence of a novel Esr1-cre mouse. Second, anxiety-like behaviors were evaluated using the elevated zero maze as previously described (44). Experimental mice were individually run on the maze for a period of 5 minutes. Total distance moved along the maze and time spent in the open arms of the maze were assessed by blinded reviewers using recorded and digitally analyzed footage via Noldus Ethovision XT 13 software (Leesburg, VA, USA).

### Hypertension Induction and Mean Arterial Pressure Recording

Following a 10-day recovery period after RSDS, induction of hypertension was performed by implantation of subcutaneous osmotic minipumps (Alzet #1002, Durect Corporation, Cupertino, CA) delivering AngII (200 ng/kg/min; Sigma #A9525, St. Louis, MO) for 2 weeks (45). The dose of AngII was chosen as a low or sub-pressor dose in control animals (46), which allowed for the assessment of sensitization after RSDS. Telemetric recording of mouse hemodynamics was performed as previously described (45). Briefly, blood pressure recordings were performed using intra-carotid arterial catheters (HD-X11, Data Sciences International, Minneapolis, MN) attached to radio telemeters for direct measurement of mean arterial pressure and heart rate in conscious unrestrained animals. Hemodynamic recordings were performed for 20 seconds every minute for the same 2 hour time periods daily (i.e., 8-10am, 12-2pm, 2-4pm) for the duration of the experiment. Averages of mean arterial pressure were calculated daily over the total 6 hour time periods when the mice displayed minimal activity and accurate resting blood pressures.

### IL-17A Quantification

At the completion of the AngII infusion, mice were sacrificed and blood collected via cardiac puncture. Plasma was isolated via centrifugation, and circulating levels of IL-17A were measured using a custom Mesoscale Discovery U-Plex kits (Mesoscale Discovery, Rockville, MD, USA) per manufacturer’s instructions. Analysis was performed using a Mesoscale QuickPlex SQ 120 and analyzed using Mesoscale Discovery software.

### Statistical Analyses

All data are presented as mean ± standard error of the mean (SEM) with sample numbers displayed as individual data points and N values are included within figure legends where appropriate. Data were first assessed for normality utilizing the Shapiro-Wilk test. Following this, data were analyzed using 2-way ANOVA and Pearson correlation where appropriate (specific tests identified in the figure legend). All statistics were calculated using GraphPad Prism (V10, GraphPad).

## 3 Results

### T-lymphocytes do not impact pro-social or anxiety-like behavior

To assess the effectiveness of the RSDS paradigm, mice were evaluated for pro-social and anxiety-like behaviors. As we have previously demonstrated (11, 44, 47-49), RSDS effectively reduced both total distanced moved and time spent in the open arms on the elevated zero maze (indicative of increased anxiety-like behavior) as well as decreased the social interaction ratio on the social interaction test (indicative of decreased pro-social behavior) in male wild-type mice (**Figure 1A-C**). Female wild-type mice demonstrated identical behavioral parameters after RSDS, demonstrating the model successfully is able to socially defeat females (**Figure 1A-C**). Intriguingly, while we have shown that T-lymphocytes are highly impacted by the effects of RSDS (11, 44, 47-49), Rag2^-/-^ mice that lack T-lymphocytes demonstrated no differences in behavior compared to wild-type mice (either male or female) after RSDS (**Figure 1D-F**). Together, these data demonstrate the RSDS paradigm is highly effective in inducing anxiety-like behavior while minimizing pro-social behavior in mice independent of T-lymphocytes.

**Figure 1.**
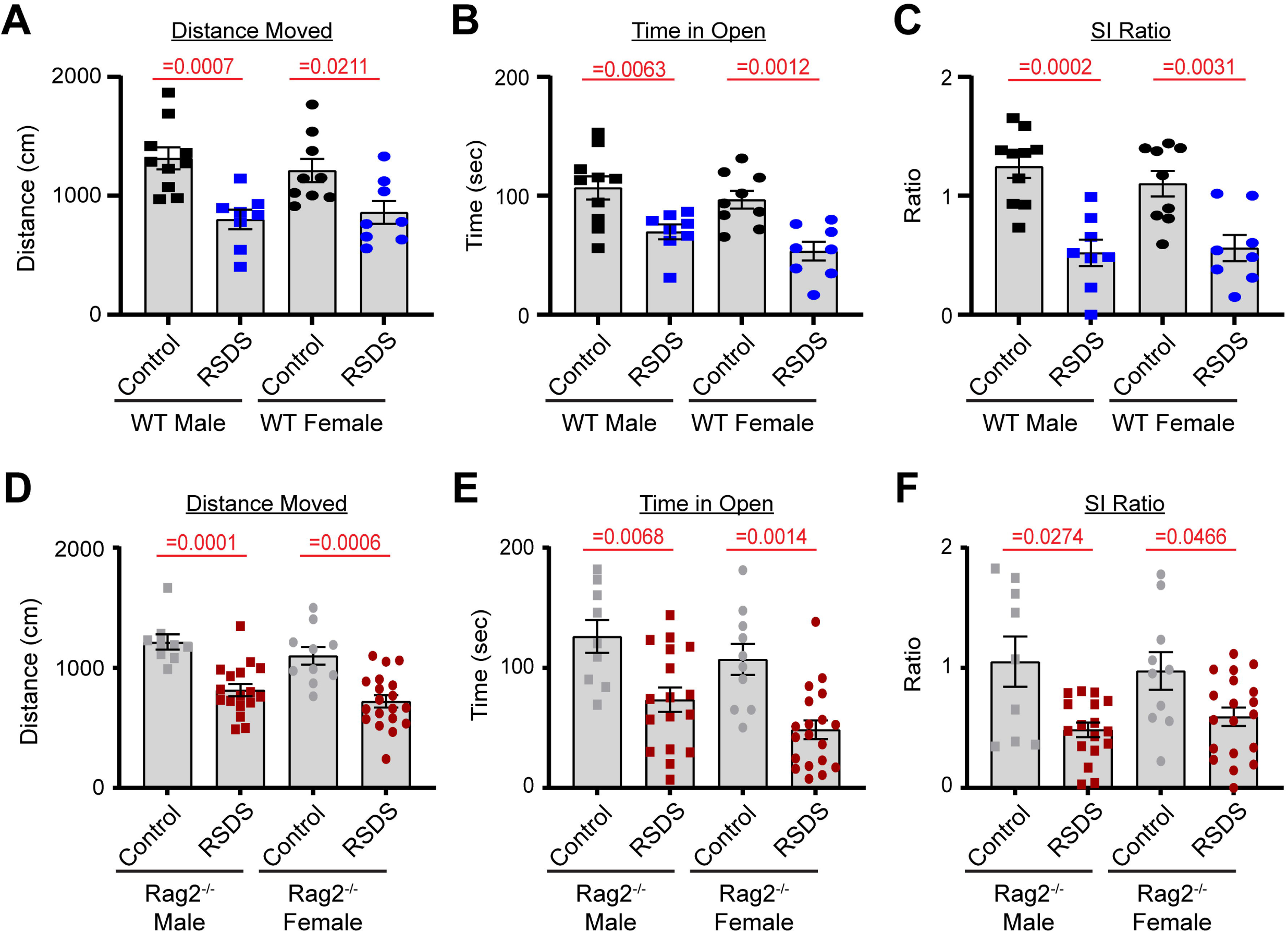
T-lymphocytes do not impact pro-social or anxiety-like behavior. RSDS was induced in male and female wild-type (WT) and Rag2^-/-^ mice followed by the assessment of pro-social behavior by the social interaction test and anxiety-like behavior by the elevated zero maze. **A**. Total distance moved on the elevated zero maze for WT mice. **B**. Time spent in the open arms of the elevated zero maze for WT mice. **C**. Social interaction (SI) ratio of WT mice in the social interaction test. **D**. Total distance moved on the elevated zero maze for Rag2^-/-^ mice. **E**. Time spent in the open arms of the elevated zero maze for Rag2^-/-^ mice. **F**. Social interaction (SI) ratio of Rag2^-/-^ mice in the social interaction test. Statistics by 2-way ANOVA with Bonferroni post-hoc.

### RSDS sensitizes mice to AngII via T-lymphocytes

Implanted radiotelemetry allows for the accurate measurement of hemodynamic parameters in conscious unrestrained mice, which limits unnecessary stress and makes it the ideal method for assessing cardiovascular pathology as a result of RSDS. As expected, wild-type mice (males and females pooled due to similar responses) showed a robust and significant elevation in mean arterial pressure during the RSDS paradigm (**Figure 2A**). However, after beginning the recovery period following RSDS, mean arterial pressure immediately returned back to baseline (**Figure 2A**). Given that we have previously demonstrated that at this same time point wild-type RSDS mice possess significantly elevated levels of systemic pro-inflammatory cytokines (11, 44, 49), we postulated that while not sufficient to raise basal blood pressure, this inflammation may sensitize mice to a hypertensive challenge such as AngII. Indeed, by infusing a subpressor dose of AngII into mice 10 days following RSDS, wild-type mice that were previously stressed demonstrated a more significant elevation in blood pressure compared to wild-type controls (**Figure 2A**). Conversely, Rag2^-/-^ mice showed no sensitization to AngII after RSDS (**Figure 2B**), even though they experienced the initial blood pressure increase during the RSDS paradigm (data not shown), which suggests T-lymphocytes mediate the RSDS-induced blood pressure sensitization to AngII.

**Figure 2.**
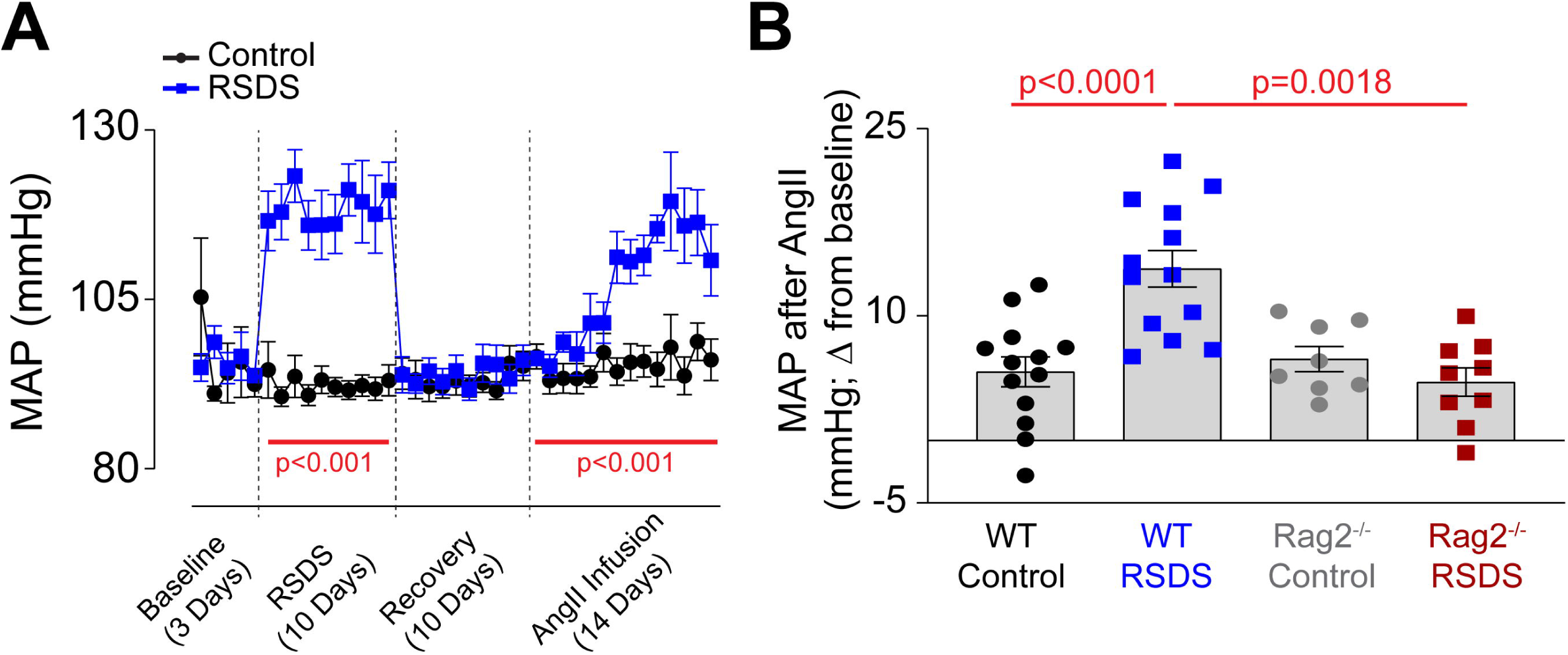
RSDS sensitizes mice to AngII via T-lymphocytes. RSDS was induced in male and female wild-type (WT) and Rag2^-/-^ mice with implanted radiotelemetry transmitters, and mean arterial pressure (MAP) was assessed daily throughout the duration of the experiment. **A**. MAP over time for the duration of the experimental paradigm in WT mice. **B**. Quantification of maximum blood pressure change from baseline to maximum level during AngII infusion in WT and Rag2^-/-^ mice. No sex differences were noted between males and females, so data are shown as pooled. Statistics by 2-way ANOVA with Bonferroni post-hoc.

### IL-17A is elevated after RSDS, but not in Rag2^-/-^ mice

The lack of an AngII sensitization response in T-lymphocyte deficient Rag2^-/-^ mice suggested an inflammatory effector molecule from these cells may be the intermediate linking the psychological trauma to the cardiovascular pathology. We have previously performed a large scale screen of inflammatory proteins that are elevated after RSDS in wild-type mice, and out of more than 40 investigated, only 8 were consistently and significantly elevated systemically after RSDS (11). However, when comparing these inflammatory proteins from wild-type mice to Rag2^-/-^ mice after RSDS, only 1 differed between the two genotypes: IL-17A (**Figure 3A**). All other 7 inflammatory proteins in Rag2^-/-^ mice showed similar levels of elevation to wild-type mice after RSDS (data not shown), suggesting they do not explain the absence of an AngII sensitization response in the Rag2^-/-^ mice. Further, both male and female wild-type mice demonstrate similarly elevated levels of IL-17A after RSDS (**Figure 3B**), which aligns with the earlier observation of a similar AngII sensitization between the sexes (**Figure 2A**). Given that IL-17A has been demonstrated to be tightly linked to the development of cardiovascular diseases (36-41), these data imply the possibility of IL-17A mediating the cardiovascular sequelae of RSDS.

**Figure 3.**
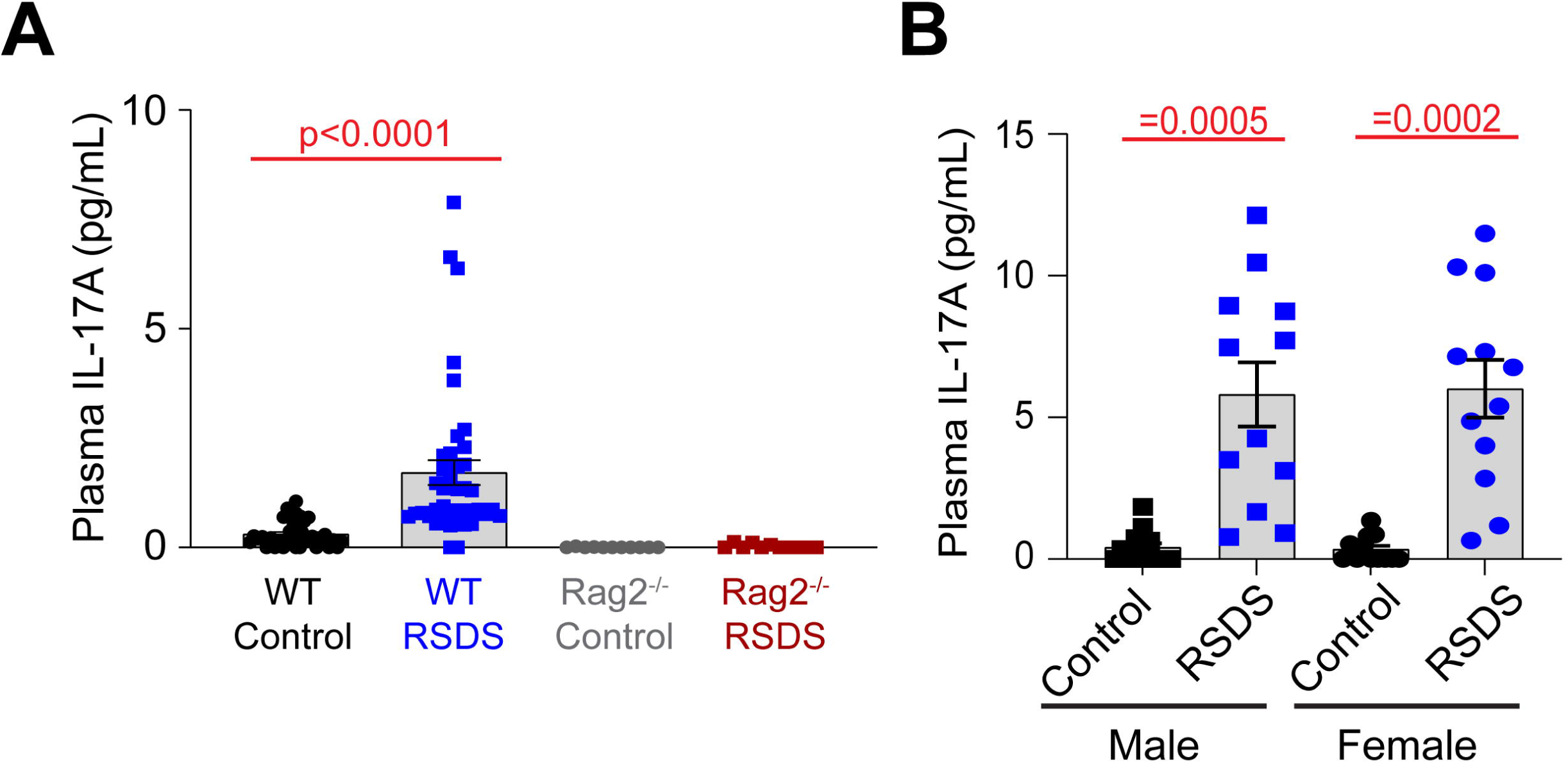
IL-17A is elevated after RSDS, but not in Rag2^-/-^ mice. RSDS was induced in male and female wild-type (WT) and Rag2^-/-^ mice followed by the extraction of peripheral blood and plasma isolation. **A**. IL-17A levels assessed by Mesoscale Discovery assay in WT and Rag2^-/-^ mice. Males and females pooled among genotypes. **B**. IL-17A levels assessed by Mesoscale Discovery assay in WT male and female animals. Statistics by 2-way ANOVA with Bonferroni post-hoc.

### AngII sensitization is linked to IL-17A, but not behavior after RSDS

To assess how the cardiovascular, inflammatory, and behavioral consequences of RSDS are related, we performed intra-animal correlations of these specific parameters. First, we found both IL-17A and AngII sensitization were positively correlated (**Figure 4A**), which aligned with previous data demonstrating the necessity of T-lymphocytes to the RSDS-induced AngII sensitization (**Figure 2B**). However, AngII sensitization showed no correlation with pro-social or anxiety-like behaviors (**Figure 4B-C**). Given the tight relationship between IL-17A and AngII sensitization, IL-17A also did not correlate with pro-social or anxiety-like behaviors (data not shown). Together, these data are highly suggestive that RSDS-induced behavioral phenotypes are not tightly coupled with pathophysiological outcomes, and that separate mechanisms may be at play within individual animals regulating separate pathways.

**Figure 4.**
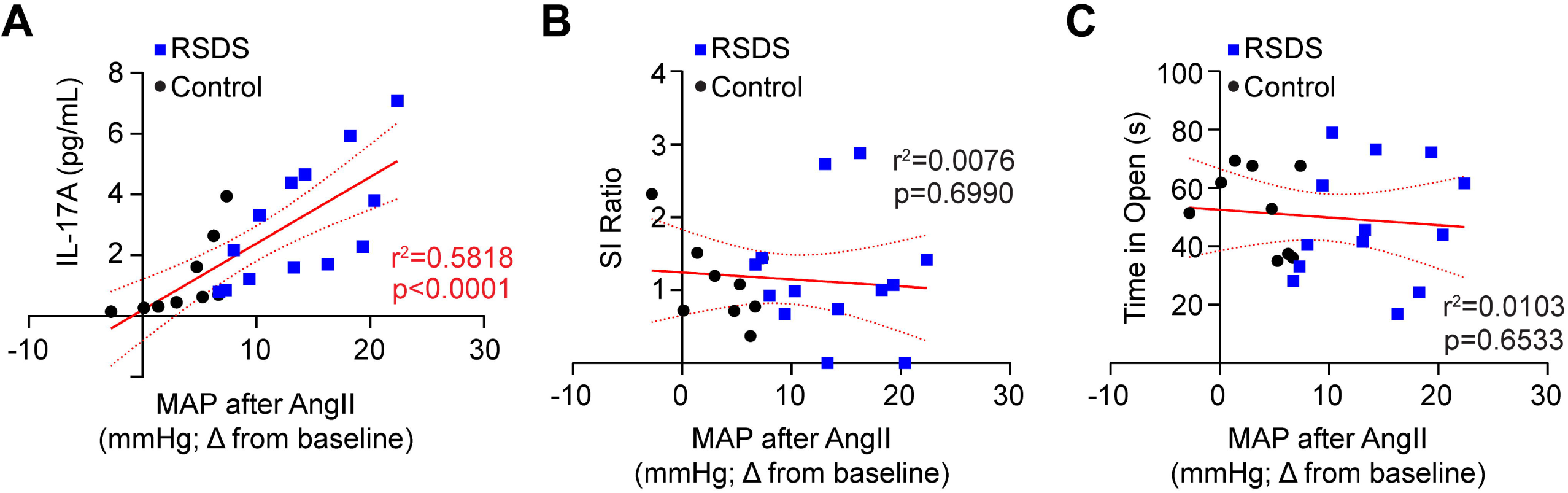
AngII sensitization is linked to IL-17A, but not behavior after RSDS. RSDS was induced in male and female wild-type (WT) with implanted radiotelemetry followed by the assessment of behavior and circulating IL-17A levels. **A**. Correlation between IL-17A and maximum mean arterial pressure (MAP) after AngII infusion. **B**. Correlation between social interaction (SI) ratio and maximum mean arterial pressure (MAP) after AngII infusion. **C**. Correlation between time spent in the open arms of the elevated zero maze and maximum mean arterial pressure (MAP) after AngII infusion. Males and females pooled. Statistics by Pearson correlation analysis.

## 4 Discussion

Herein, we explored the relationship between psychological trauma, immune dysregulation, and cardiovascular dysfunction. First, we found that using chemogenetically-altered aggressive mice was sufficient to induce social defeat behavior in both males and females with no apparent sex differences observed in the parameters assessed. Second, we detected that T-lymphocytes do not appear directly involved in the behavioral consequences of RSDS, as T-lymphocyte deficient Rag2^-/-^ mice displayed identical behavioral patterns to wild-type mice in both males and females. Third, RSDS is sufficient to produce a blood pressure sensitization to exogenous AngII, which is dependent upon T-lymphocytes and tightly linked to systemic IL-17A levels. Last, RSDS-induced AngII sensitization was not associated with the behavioral consequences of RSDS, suggesting the potential for disparate mechanisms underlying psychopathology versus pathophysiology. Together, our results demonstrate a new brain-immune-cardiovascular axis that is impacted by psychological trauma.

Over the last few decades, numerous meta-analyses have demonstrated tight associations between PTSD and the development of cardiovascular disease (15-20). While incredibly informative in elucidating these links, these studies have also raised many questions. For example, given the results of the studies, many investigators often presume that PTSD causes cardiovascular disease, but it is just as likely that cardiovascular disease (or at least some underlying pathophysiology related to cardiovascular disease, such as inflammation) elevates the risk of developing PTSD after psychological trauma. While this directional quandary has been excellently discussed and evaluated (50), the take home is that the majority of clinical studies have not been designed to address this specific question, though some have provided evidence suggesting a bidirectionality exists. For example, by using a longitudinal study design of pre-versus post-deployment military personnel, several studies have uncovered certain inflammatory signatures that existed prior to the development of PTSD after psychological trauma (51-56). These again do not specifically delineate cause and effect, but suggest that the presence of specific elevated inflammatory markers may predict those who develop PTSD. Conversely, several studies have identified that PTSD does appear to increase inflammation over time (57-60), which may underlie the elevated risk for cardiovascular disease. While these conclusions may seem at odds, it is highly possible that both situations are occurring simultaneously. In other words, individuals with elevated inflammation possibly have a lower threshold or decreased resilience to psychological trauma, which predisposes them to PTSD development, but trauma and PTSD further elevate the inflammation pushing the body into somatic pathophysiology, such as cardiovascular disease. Our work using rodents supports the notion that psychological trauma elevates inflammation (11, 42, 44, 48, 49, 61, 62), though we have not comprehensively explored inflammatory parameters pre-RSDS and how they impact chronic outcomes. Importantly, given our data presented herein which suggest behavior and physiology may not be tightly coupled, it would be imperative to explore not only how inflammation pre-RSDS shapes behavioral outcomes after psychological trauma, but also consequences that are physiological in nature.

While it may be intuitive to assume that psychopathology and pathophysiology are tightly linked following psychological trauma, this concept appears much more nuanced than a simple binary relationship. For example, if PTSD patients develop inflammation or cardiovascular disease, then by definition psychopathology and pathophysiology will be coupled in these individuals given that a diagnosis of PTSD requires a specific subset of behavioral pathology. However, not all PTSD patients develop inflammation or cardiovascular diseases, and emerging evidence also suggests that exposure to psychological trauma alone may be sufficient to generate pathophysiology in the absence of psychopathology (25). These investigations have demonstrated that traumatic stress alone was sufficient to increase circulating inflammation (63, 64), elevate mitochondrial dysfunction (65, 66), and alter the function of the renin-angiotensin system (67, 68). Our data presented herein, as well as our previously published observations (11), also demonstrate that exposure to traumatic stress elevates inflammation and cardiovascular pathology independently of behavioral manifestations. Together, these findings suggest that traumatic stress can produce multiple phenotypes: individuals with psychopathology and pathophysiology, those with one or the other, or those with neither. These phenotypes may be even more subcategorized when taking into consideration the specific dimensions of behavioral changes as well as the particular pathophysiology. Importantly, these findings have significant clinical implications in that currently exposure to psychological trauma is not an actively treated “disease.” Only once that exposure develops into a form of chronic psychopathology is the patient therapeutically managed, which implies if no over behavioral pathology develops, the patient is essentially considered “healthy” with no additional consideration of their potential underlying pathophysiology. These current unknowns warrant longitudinal studies exploring these specific outcomes of psychological trauma and how behavior and physiology are related.

The study presented here does possess certain limitations. First, while we have identified an association between IL-17A and the blood pressure sensitization to AngII, we have not yet mechanistically defined this relationship. In future studies, we intend on using both pharmacological (i.e., neutralizing antibodies) and genetic methods of reducing IL-17A levels in RSDS to assess the chronic cardiovascular outcomes. Second, this study only explored two dimensions of behavior: pro-social and anxiety-like. While there were no direct associations with these behaviors and the RSDS-induced pathophysiology, it is highly possible that these specific psychopathologies are not coupled with these particular pathophysiologies. The exploration of more behaviors such as anhedonia, depression-like behavior, cognition, and many others are highly warranted to truly uncover any nuance linking behavior to physiology. Third, while we have uncovered that RSDS-induces an increased pressor response to AngII, it remains unknown if other cardiovascular challenges (e.g., high salt, nitric oxide sequestration, coronary ligation, etc.) also demonstrate sensitization. Last, while RSDS is an accepted as a model of psychological trauma, it does possess reports of non-PTSD outcomes such as decreased fear extinction and increased HPA feedback (69). Validating the results presented herein using additional models of psychological trauma will aid in the understanding of the broader applicability of our findings.

In summary, this investigation has uncovered an apparent pathway linking psychological trauma to cardiovascular dysfunction via T-lymphocyte-mediated inflammation. Given the lack of a correlation between the psychopathology and pathophysiology in this model, our data suggests that psychological trauma alone is sufficient to induce pathophysiology, which warrants deeper exploration into patients exposed to traumatic stress independent of their development of psychopathology. In doing so, clinical practice may shift to a model of intervening as soon as possible to the traumatic event in hopes of preventing downstream negative behavioral and physiological consequences.

## 6 Author Contributions

AJC designed research studies. All authors conducted experiments, acquired data, and/or performed analyses. AJC wrote the manuscript, while all authors approved the final version of the manuscript. AJC provided funding and experimental oversight.

## 8 Data Availability Statement

All data used in this manuscript are available upon request to interested researchers.

